# HAPDeNovo: a haplotype-based approach for filtering and phasing *de novo* mutations in linked read sequencing data

**DOI:** 10.1101/220830

**Authors:** Xin Zhou, Serafim Batzoglou, Arend Sidow, Lu Zhang

## Abstract

**Background:** *De novo* mutations (DNMs) are associated with neurodevelopmental and congenital diseases, and their detection can contribute to understanding disease pathogenicity. However, accurate detection is challenging because of their small number relative to the genome-wide false positives in next generation sequencing (NGS) data. Software such as DeNovoGear and TrioDeNovo have been developed to detect DNMs, but at good sensitivity they still produce many false positive calls.

**Results:** To address this challenge, we develop HAPDeNovo, a program that leverages phasing information from linked read sequencing, to remove false positive DNMs from candidate lists generated by DNM-detection tools. Short reads from each phasing block are allocated to each of the two haplotypes followed by generating a haploid genotype for each putative DNM.HAPDeNovo removes variants that are called as heterozygous in one of the haplotypes because they are almost certainly false positives. Our experiments on 10X Chromium linked read sequencing trio data reveal that HAPDeNovo eliminates 80% to 99% of false positives regardless of how large the candidate DNM set is.

**Conclusions:** HAPDeNovo leverages the haplotype information from linked read sequencing to remove spurious false positive DNMs effectively, and it increases accuracy of DNM detection dramatically without sacrificing sensitivity.

## Background

*De novo* mutations (DNMs) have been shown to be a major cause of neurodevelopmental and other congenital diseases including autism [1], schizophrenia [2], intellectual disability [3], and congenital heart disease [4]. Next generation sequencing of nuclear families provides an unprecedented opportunity to investigate the *de novo* mutation spectrum of these diseases at single nucleotide resolution. Variant calling methods such as GATK [5] and SAMtools [6] implement a straightforward approach to explore DNMs by selecting the mutations that appear in the child but not the parents. Other approaches such as DeNovoGear [7] and PolyMutt [8] model the family relationship as a prior probability of mutation transmission to distinguish true DNMs from noise, dramatically improving performance. These programs assume a consistent mutation rate across all positions, which is not always the case. TrioDeNovo [9] was developed to address this issue by employing flexible priors. Nevertheless, most of the existing algorithms are still overwhelmed by an enormous number of false positives, which are probably caused by factors such as sequencing coverage bias, sequencing batch effects, and alignment artifacts on repetitive regions. There has been a lack of studies that investigate which factor has the most impact and how to correct biases in *de novo* mutation calling. Phasing of inherited variants, inferred by linkage disequilibrium or allele transmission, has commonly been applied to refine inherited variant calls [10–12]. Knowing the phase of DNMs is critical in determining their parent-of-origin. Yet, phasing is complicated and remains challenging for DNMs on just short reads generated from the typical ~500bp fragments in Illumina sequencing. The 10X Chromium system microfluidically partitions long DNA fragments from which short fragments and, subsequently, Illumina reads are generated [13]. Thus, each original long DNA fragment generates a collection of short reads with a shared barcode (linked reads), enabling robust and accurate, genome-scale variant genotyping and phasing. Phasing analysis reveals that linked read sequencing generates a very low overall long switch error (<0.03%) [14]. Here we developed a novel filtering and phasing toolkit for DNMs, HAPDeNovo, which takes full advantage of robust variant phasing from linked read sequencing to sift true DNMs from noise. We show that HAPDeNovo drastically eliminates false positive DNMs without decreasing the detection rate of true positives. We identify the culprit of false positive calls to be allele-specific sequencing coverage biases.

## Implementation

Linked read sequencing is a technology that allows for simultaneous variant calling and phasing, by reconstructing the original long fragments from linked short reads. When reads with the same barcode align in proximity to each other in the genome, they originated from the same haplotype because the original template was a single DNA fragment. In general, HAPDeNovo is designed to re-calibrate the DNM quality based on read coverage and sequencing quality for each haplotype. The reads from each phasing block are allocated to either of the two haplotypes, enabling HAPDeNovo to identify two haploid genotypes for each candidate DNM. Each putative *de novo* mutation (homozygous reference allele for both parents and heterozygosity for the child) is categorized as high-confidence if all the three genotypes in the trio can be phased and all haploid genotype calls are homozygous. The remaining DNMs would be categorized to be low-confidence if there are no reads covering one or more haplotypes, since there is insufficient information to ascertain whether they are false positives. HAPDeNovo involves three steps: 1. Variant calling and phasing; 2. Haplotype-specific genotyping; 3. Removing false positive DNMs.

The input to HAPDeNovo is paired-end reads in FASTQ files generated by Illumina-sequencing of 10X Chromium libraries for each individual of the trio. In the first step, all the reads are aligned to the reference genome by LongRanger or another barcode-aware aligner and HAPDeNovo would perform multi-sample variant calling on the trio with any available variant callers (FreeBayes by default). Because variant phasing is independent for each individual, HAPDeNovo separates the variants into three individual VCF files for variant phasing based on the barcode-aware haplotype assembly approaches such as Long Ranger or HapCUT2 [15]. The individual phased VCF files are then merged into a phased multi-sample VCF file. For each phase block, HAPDeNovo determines the haplotype that each read comes from and marks the read accordingly in the BAM file.

In the second step, the BAM file for each individual is divided into three, according to the three haplotype tags: HP1, HP2 and HP0, which denote the reads coming from maternal haplotype, paternal haplotype, and undetermined haplotype within the phase block, respectively. Then, multi-sample variant calling is performed again on all nine BAM files, which identifies the specific alleles that comprise each individual’s two haplotypes.

In the last step, putative DNMs are extracted into a VCF (Variant Call Format) file from the original multi-sample variant calls (FreeBayes by default), if the allele occurs in only one haplotype of the child (heterozygous variant) and is absent in both parents (homozygous reference). HAPDeNovo currently accepts multi-sample variant calls to produce the DNM candidate set from GATK, TrioDeNovo and DeNovoGear, for which preprocessing scripts are included in HAPDeNovo. HAPDeNovo defines variant sites to be high-confidence if all the six genotypes called from the reads with HP1 and HP2 tags are homozygous (0|0 or 1|1, denoting reference and variant haploid genotypes) and the calls are supported by sufficient sequencing depth (>1X by default). High-confidence DNMs are defined as such when they belong to the high-confidence sites, all four parental haploid alleles are 0, the child’s alleles are 0 and 1, and the DNM is phased. The candidate DNMs are defined as low-confidence when one or more haplotype is uncovered by any reads, but they are identified as putative DNMs in the original candidate set. These low-confidence variants are kept for further consideration since HAPDeNovo is unable to determine on the basis of the haploid genotypes whether they are false positives.

It is a special case that variants on the male’s X chromosome are naturally phased for the non-pseudoautosomal regions, so their genotypes are directly translated to haplotype calls. The variants from pseudoautosomal regions are merged into autosomal chromosomes for analysis; in practice, mapping quality is low in the pseudoautosomal regions because they are duplicated in the reference genome. If the child is female, putative *de novo* mutations are categorized as high-confidence if the two genotypes from the mother and child can be phased and all five haploid calls (including father’s X chromosome) are homozygous. If the child is male, only the haploid calls on the X chromosomes of mother and child are considered.

The required sequencing depth per haplotype is a user-defined parameter. The output of HAPDeNovo is a flat file containing high-confidence DNMs annotated by *H*, and low-confidence DNMs annotated by *L*.

## Results

The performance of HAPDeNovo was evaluated on the validated DNMs of the 1000 Genomes Project CEU trio (NA12878, daughter; NA12891, father; and NA12892, mother). We downloaded the reads of the three samples that were generated by the 10X Chromium system from the Genome In A Bottle website (See **Availability of data and materials**). All of them have sufficient sequencing depth for variant calling (NA12878: 300Gb reads, 74.96X coverage; NA12891: 128Gb reads, 36.94X coverage; NA12892: 128Gb reads, 36.88X coverage). There are 49 validated germline *de novo* mutations in NA12878 [16], serving as a gold standard (Additional file 5: Table S5). The alternative alleles of 45 of these were covered by 10X-based linked read sequencing at least once; four *de novo* mutations could not be evaluated due to poor sequencing coverage of alternative alleles (Supplementary Table S5).

We used Lariat [17] to align the reads from the trio against the reference genome (hg19) followed by variant calling and generation of a set of putative DNMs. More than 95% of the reads can be aligned to the reference genome (NA12878: 96.17%, NA12891: 97.15%, NA12892: 96.86%). We included the variant calls from four programs: two general purpose callers (GATK and FreeBayes) and two DNM specific callers (TrioDeNovo and DeNovoGear) to evaluate the impact of different inputs with respect to HAPDeNovo performance. To incorporate as many DNMs in the gold standard as possible, we applied lenient parameters in variant calling. A depth threshold was commonly applied to the variants from all four methods as well as additional unique threshold for each program to pre-filter those false positives before phasing. De Novo Quality (DQ) and Posterior Probability (PP) were considered in TrioDeNovo and DeNovoGear, and Genotype Likelihoods (GL or PL) were applied to GATK and FreeBayes. We also varied these thresholds to examine their potential influence (Additional file 1: Table S1-Additional file 4: Table S4).

If only DNMs with sequencing depth greater than 15X were taken into consideration, a majority of the variant sites were classified as high-confidence (85.3% for FreeBayes and TrioDeNovo, 99.5% for GATK, and 98.9% for DeNovoGear). FreeBayes initially generated 10,431 candidate DNMs but after application of a GL filter (GL= −50), 5,829 candidates remained, including all 44 true positives. TrioDeNovo generated 3,955 candidate DNMs; after application of a more stringent quality threshold (DQ=7), 3,717 candidates remained, including all 44 true positives. GATK and DeNovoGear generated much larger candidate sets (242,530 and 89230 after applying PL=450 and PP=3E-5, respectively), and each missed one true DNM (Table 1).

**Table 1:** The performance of FreeBayes, TrioDeNovo, GATK, and DeNovoGear with sequencing depth threshold 15 and their optimal parameters (GL = −50 for FreeBayes, DQ = 7 for TrioDeNovo, PL = 450 for GATK, and PP = 1E-4 for DeNovoGear). After further applying HAPDeNovo, the number of false positives decreases significantly for all four inputs. HAPDenovo also calculates the confidence of DNMs. A high proportion of TP (33/44, 32/43) comes from high-confidence DNMs. **TP** (True Positive): number of *de novo* mutations in both candidate set and the gold standard. **FP** (False Positive): number of mutations in the candidate set but not in the gold standard. **HC** (High Confidence): high-confidence DNMs. **LC** (Low Confidence): low-confidence DNMs.

|  | Optimal-Paras |  | HAPDeNovo |  | HAPDeNovo |  |  |  |
| --- | --- | --- | --- | --- | --- | --- | --- | --- |
|  |  |  |  |  | HC |  | LC |  |
|  | TP | FP | TP | FP | TP | FP | TP | FP |
| **FreeBayes** | 44 | 5785 | 44 | 1083 | 33 | 219 | 11 | 864 |
| **TrioDeNovo** | 44 | 3673 | 44 | 674 | 33 | 124 | 11 | 550 |
| **GATK** | 43 | 242487 | 43 | 1396 | 32 | 258 | 11 | 1138 |
| **DeNovoGear** | 43 | 89187 | 43 | 1241 | 32 | 246 | 11 | 995 |

Application of HAPDeNovo to these call sets eliminated a considerable number of false positives (81.3% for FreeBayes, 81.6% for TrioDeNovo, 99.4% for GATK, 98.6% for DeNovoGear; Table 1), without sacrificing any true positives. The number of remaining candidate DNMs was similar among the inputs (1,439 for GATK, 1,127 for FreeBayes, 1,284 for DeNovoGear, 718 for TrioDeNovo; Table 1). In each of these sets, HAPDeNovo identified approximately 22% of as high-confidence DNMs, which included a majority of true positives (33/44 for FreeBayes and TrioDeNovo and 32/43 for GATK and DeNovoGear). By increasing the stringency of thresholds, further false positive reduction was achieved at a small cost of sensitivity (Additional file 1: Table S1-Additional file 4: Table S4). Moreover, HAPDeNovo could phase and determine the parent-of-origin of all the 44 validated *de novo* mutations.

To understand whether the increased specificity of HAPDeNovo is sensitive to read depth of raw variant calls from the four programs, we performed an extensive ROC analysis for each of the four variant callers with and without HAPDeNovo (Figure 2). We applied the read depth thresholds from 10X to 30X to the raw variants and found the optimal parameter settings for the four variant callers (the maximal number of true DNMs and minimal number of false positives; GL=-50 and PP=3E-5 for FreeBayes and DeNovoGear, DQ=7 and PL=450 for TrioDeNovo and GATK). Application of HAPDeNovo on top of the optimal parameter settings always generated a smallest set of false positives without losing any true positives. In general, 80% to 99% of false positives were eliminated by HAPDeNovo. Specifically, by using HAPDeNovo, the average false positive removal was 82.7% for FreeBayes, 82.7% for TrioDeNovo, 99.5% for GATK, and 98.8% for DeNovoGear.

**Figure 1:**
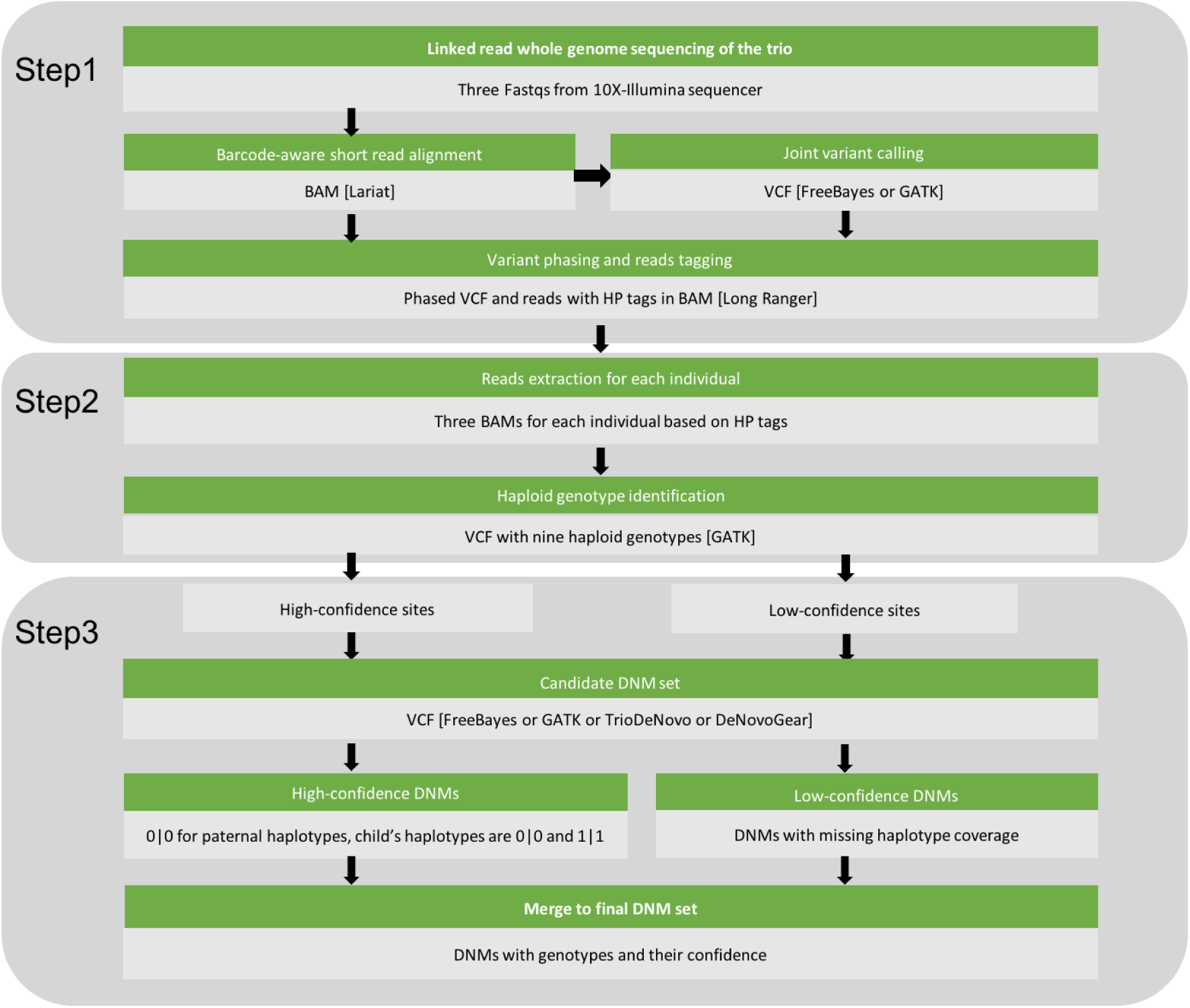
Pipeline of HAPDeNovo. Software is in brackets. HP: haplotype

**Figure 2:**
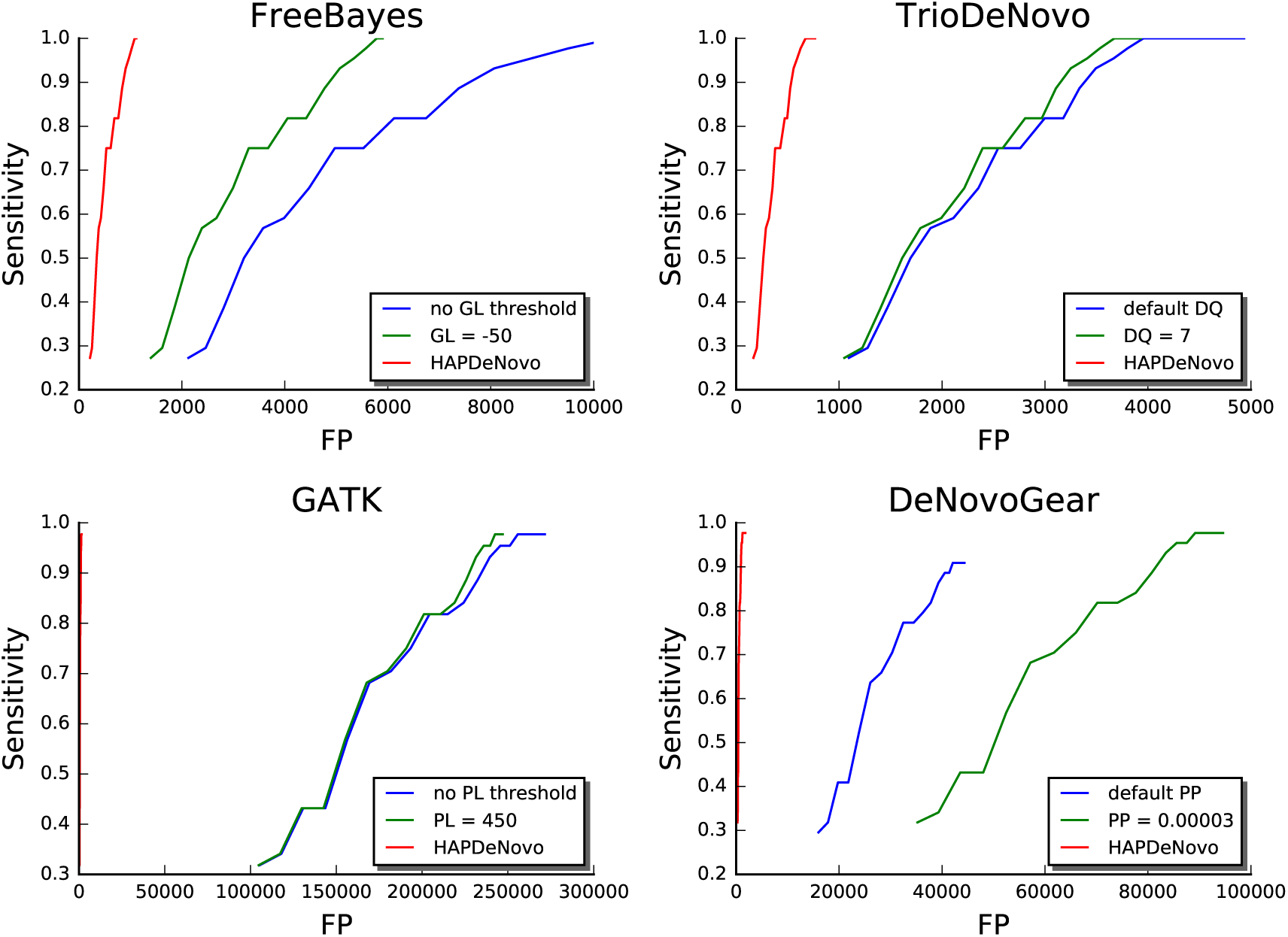
ROC curves of DNMs called by FreeBayes, TrioDeNovo, GATK, and DeNovoGear by optimal parameter setting, and the improved ROC curves after applying HAPDeNovo (red line). Sequencing depth threshold is varied from 10 (start of each plot line, leftmost point) to 30 (end of each plot line, rightmost point). **FP** (False Positive): Number of false positive DNMs. **Sensitivity**: Number of true positive DNMs divided by the total number of true positive plus false negative DNMs. GL: Genotype Likelihood; DQ: De Novo Quality; PL: Posterior Likelihood; PP: Posterior Probability. **Blue curves** show the sensitivity and number of FPs at default settings (no GL and no PL thresholds for FreeBayes and GATK, respectively). **Green curves** show the sensitivity and number of FP at optimal parameter settings (GL = −50 for FreeBayes, DQ = 7 for TrioDeNovo, PL = 450 for GATK, and PP = 3E-5 for DeNovoGear). **Red curves** show the performance after applying HAPDeNovo.

To ascertain whether haplotype information was generally beneficial for calling DNMs we also analyzed results from Long Ranger, which, like HAPDeNovo, can allocate allele-specific reads to each haplotype. This boosts the power for detecting heterozygous variants, such as DNMs. We compared the performance of TrioDeNovo, Long Ranger and HAPDeNovo with respect to DNM calling. Both Long Ranger and HAPDeNovo performed better than TrioDeNovo, which is consistent with the idea that the accuracy of calling DNMs benefits from the haplotype information. Nevertheless, Long Ranger, which considers the individuals of a trio independently from one another, called many more false positives than HAPDeNovo. HAPDeNovo eliminated ~80% (Table 2) of false positives from Long Ranger, suggesting that HAPDeNovo's simultaneous consideration of all six haplotypes boosts accuracy in DNM detection.

**Table 2:** Comparing the performance of TrioDeNovo, Long Ranger and HAPDeNovo (using Tri-oDeNovo as input) as a function of sequencing depth ranging from 10 to 20. DQ = 7 was used as the quality threshold for TrioDeNovo. **TP** (True Positive): Number of DNMs in candidate set plus gold standard. **FP** (False Positive): Number of DNMs in the candidate set but not in the gold standard.

|  | Depth | 10 | 11 | 12 | 13 | 14 | 15 | 16 | 17 | 18 | 19 | 20 |
| --- | --- | --- | --- | --- | --- | --- | --- | --- | --- | --- | --- | --- |
| **TrioDeNovo** | **TP** | 44 | 44 | 44 | 44 | 44 | 44 | 43 | 42 | 41 | 39 | 36 |
|  | **FP** | 3932 | 3926 | 3923 | 3862 | 3789 | 3673 | 3532 | 3410 | 3250 | 3106 | 2969 |
| **Long Ranger** | **TP** | 44 | 44 | 44 | 44 | 44 | 44 | 43 | 42 | 41 | 39 | 36 |
|  | **FP** | 3911 | 3907 | 3904 | 3844 | 3771 | 3655 | 3515 | 3394 | 3235 | 3092 | 2956 |
| **HAPDeNovo** | **TP** | 44 | 44 | 44 | 44 | 44 | 44 | 43 | 42 | 41 | 39 | 36 |
|  | **FP** | 768 | 766 | 765 | 744 | 715 | 674 | 626 | 593 | 558 | 525 | 496 |

Finally, we explored whether HAPDeNovo’s consideration of reads that cannot be allocated to a specific haplotype (HP0) would affect the accuracy of DNM calling. We compared HAPDeNovo performance when only HP1 and HP2 BAM files are provided as input versus when all nine BAM files (including those of HP0) were considered. Accuracy without HP0 is lower than with HP0 (Table 3).

**Table 3:** Comparing the performance of HAPDeNovo using HP1 and HP2 only versus HP0, HP1, and HP2 (both with TrioDeNovo as input), with DQ=7 and sequencing depth changing from 10 to 20. **TP** (True Positive): Number of DNMs in candidate plus gold standard. **FP** (False Positive): Number of DNMs in the candidate set but not in the gold standard.

|  | Depth | 10 | 11 | 12 | 13 | 14 | 15 | 16 | 17 | 18 | 19 | 20 |
| --- | --- | --- | --- | --- | --- | --- | --- | --- | --- | --- | --- | --- |
| **TrioDeNovo** | **TP** | 44 | 44 | 44 | 44 | 44 | 44 | 43 | 42 | 41 | 39 | 36 |
|  | **FP** | 3932 | 3926 | 3923 | 3862 | 3789 | 3673 | 3532 | 3410 | 3250 | 3106 | 2969 |
| **HAPDeNovo**<br>**by HP1, HP2** | **TP** | 40 | 40 | 40 | 40 | 40 | 40 | 40 | 39 | 39 | 37 | 35 |
|  | **FP** | 605 | 604 | 603 | 584 | 557 | 521 | 481 | 453 | 423 | 394 | 372 |
| **HAPDeNovo**<br>**by HP1, HP2, HP0** | **TP** | 44 | 44 | 44 | 44 | 44 | 44 | 43 | 42 | 41 | 39 | 36 |
|  | **FP** | 768 | 766 | 765 | 744 | 715 | 674 | 626 | 593 | 558 | 525 | 496 |

Finally, we separately analyzed the X chromosome independently from the autosomes. There is only one validated germline *de novo* mutation on the X chromosome in NA12878, serving as a gold standard (Supplementary Table S5). For FreeBayes and TrioDeNovo, a depth threshold of 15X retained the smallest number of false positives and kept the true one. With the inputs from these two programs, HAPDeNovo decreased the false positives by 84.0% without sacrificing the true positive (Supplementary Table S6a). For GATK, the optimal depth threshold for the X chromosome was 39X, and HAPDeNovo removed 99.9% of false positives, which was similar to its removal rate for autosomes (Supplementary Table S6b). Eliminating any false positives from DeNovoGear would also lose the true one simultaneously by applying threshold for read depth. DeNovoGear removed the gold standard SNP on the X chromosome regardless of parameter settings (Supplementary Table S6c).

## Discussion

Many diseases with early onset age are associated with *de novo* mutations. The extensive availability of next generation sequencing technology has encouraged the study of *de novo* mutations, which played an important role in explaining why diseases with critically decreased fitness occur frequently in the human population [12]. Barcode-based linked read sequencing, as an alternative solution to single molecule long reads sequencing, enables high-quality haplotype phasing and structural variation analysis [14]. In this study, we developed HAPDeNovo, a flexible and efficient pipeline that benefits from variant phasing from linked read sequencing to improve *de novo* mutations calling. The assignment of reads to each haplotype (HP1, HP2) decreases the chance that a genotype is miscalled because one haplotype is dominant, such as the haplotype with the reference allele in the parents. This is the major cause of an inherited variant getting called as a *de novo* mutation.

To date, methods that were developed to work on short reads alone have not achieved satisfactory performance for calling DNMs. For example, GATK best practices is highly effective in reducing the impact of sequencing and alignment artifacts on variant calls, but it is still challenged in the accurate detection of *de novo* mutations. Existing *de novo* mutation-specific callers like DeNovoGear and TrioDeNovo perform better than the general callers such as GATK and FreeBayes. Nevertheless, tremendous numbers of false positives remain in their ultimate results. We showed HAPDeNovo to be superior in comparison because it explicitly leverages the haplotype-specific genotypes of the three individuals of a trio simultaneously.

## Conclusions

Linked read sequencing is a powerful tool to phase the variants from a single person rather than by statistical inference from a population. This boosts our ability to identify the parent-of-origin and transmission of *de novo* mutations. HAPDeNovo introduces haploid genotyping to take advantage of physical phasing that benefits from linked read sequencing and to overcome sequencing coverage imbalance and alignment artifacts in detecting *de novo* mutations. HAPDeNovo can be applied in conjunction with any variant caller to dramatically decrease false positive mutations. HAPDeNovo is user friendly and includes auxiliary scripts to process the results from other tools, and in the future, will be extended to detect inherited mutations in complex pedigrees and somatic mutations in tumor-normal pairs.

## Availability and requirements

**Project name:** HAPDeNovo.

**Project home page:** https://github.com/maiziex/HAPDeNovo.

**Operating system(s):** Linux.

**Programming language:** Python & Bash.

**Other requirements:** GATK and FreeBayes.

**License:** GNU GPLv2.

**Any restrictions to use by non-academics:** None.

## Additional files

**Additional file 1: Table S1**. Comparing the performance between TrioDeNovo and TrioDeNovo+HAPDeNovo with sequencing depth changing from 10 to 30 and different values of DQ. **TP** (True Positive): the number of DNMs mutations in both candidate set and the gold standard. **FP** (False Positive): the number of mutations belongs to the candidate set but not in the gold standard.

**Additional file 2: Table S2**. Comparing the performance between FreeBayes and FreeBayes+HAPDeNovo with sequencing depth changing from 10 to 30 and different values of GL. **TP** (True Positive): the number of DNMs in both candidate set and the gold standard. **FP** (False Positive): the number of DNMs belongs to the candidate set but not in the gold standard.

**Additional file 3: Table S3**. Comparing the performance between GATK and GATK+HAPDeNovo with sequencing depth changing from 10 to 30 and with different values of PL. **TP** (True Positive): the number of DNMs in both candidate set and the gold standard. **FP** (False Positive): the number of DNMs belongs to the candidate set but not in the gold standard.

**Additional file 4: Table S4**. Comparing the performance between DeNovoGear and DeNovoGear+HAPDeNovo with sequencing depth changing from 10 to 30 and with different values of PP. **TP** (True Positive): the number of DNMs in both candidate set and the gold standard. **FP** (False Positive): the number of DNMs belongs to the candidate set but not in the gold standard.

**Additional file 5: Table S5**. 48 *de novo* mutations for NA12878 (hg19) validated by sanger sequencing

**Additional file 6: Table S6**. Comparing the performance only on X chromosome for FreeBayes, TrioDenovo, GATK, and DeNovoGear before and after applying HAPDeNovo.

## Abbreviations

DNMs: *De novo* mutations; NGS: Next Generation Sequencing; DQ: De Novo Quality; PP: Posterior Probability; GL: Genotype Likelihoods; PL: Phred-scaled Likelihood; VCF: Variant Call Format; ROC: Receiver Operating Characteristic

Ethic approval and consent to participate

Not applicable

Consent for publication

Not applicable

## Availability of data and materials

**NA12878 dataset**

ftp://ftp-trace.ncbi.nlm.nih.gov/giab/ftp/data/NA12878/10Xgenomics_ChromiumGenome_LongRanger2.0_06202016/

**NA12891 dataset**

ftp://ftp-trace.ncbi.nlm.nih.gov/giab/ftp/technical/10Xgenomics_ChromiumGenome_LongRanger2.0_06202016/NA12891/

**NA12892 dataset**

ftp://ftp-trace.ncbi.nlm.nih.gov/giab/ftp/technical/10Xgenomics_ChromiumGenome_LongRanger2.0_06202016/NA12892/

**HAPDeNovo**

https://github.com/maiziex/HAPDeNovo

Author’s contributions

XZ and LZ wrote the software and performed the analyses. LZ, AS and SB conceived the study. XZ, LZ, SB and AS analyzed the results and wrote the paper. LZ and AS read and approved the final manuscript.

## Acknowledgements

AS acknowledges support from NIH/NCI and the BRCA foundation. XZ was supported by a training grant (NIST/JIMB).

## References

1. Ronemus M, Iossifov I, Levy D, Wigler M: The role of de novo mutations in the genetics of autism spectrum disorders. Nat Rev Genet 2014, 15(2):133–141.

2. Fromer M, Pocklington AJ, Kavanagh DH, Williams HJ, Dwyer S, Gormley P, Georgieva L, Rees E, Palta P, Ruderfer DM et al: De novo mutations in schizophrenia implicate synaptic networks. Nature 2014, 506(7487):179–+.

3. Gregor A, Oti M, Kouwenhoven EN, Hoyer J, Sticht H, Ekici AB, Kjaergaard S, Rauch A, Stunnenberg HG, Uebe S et al: De Novo Mutations in the Genome Organizer CTCF Cause Intellectual Disability. Am J Hum Genet 2013, 93(1):124–131.

4. Homsy J, Zaidi S, Shen YF, Ware JS, Samocha KE, Karczewski KJ, DePalma SR, McKean D, Wakimoto H, Gorham J et al: De novo mutations in congenital heart disease with neurodevelopmental and other congenital anomalies. Science 2015, 350(6265):1262–1266.

5. DePristo MA, Banks E, Poplin R, Garimella KV, Maguire JR, Hartl C, Philippakis AA, del Angel G, Rivas MA, Hanna M et al: A framework for variation discovery and genotyping using next-generation DNA sequencing data. Nature genetics 2011, 43(5):491–+.

6. Li H, Handsaker B, Wysoker A, Fennell T, Ruan J, Homer N, Marth G, Abecasis G, Durbin R, Proc GPD: The Sequence Alignment/Map format and SAMtools. Bioinformatics 2009, 25(16):2078–2079.

7. Ramu A, Noordam MJ, Schwartz RS, Wuster A, Hurles ME, Cartwright RA, Conrad DF: DeNovoGear: de novo indel and point mutation discovery and phasing. Nat Methods 2013, 10(10):985–+.

8. Li BS, Chen W, Zhan XW, Busonero F, Sanna S, Sidore C, Cucca F, Kang HM, Abecasis GR: A Likelihood-Based Framework for Variant Calling and De Novo Mutation Detection in Families. Plos Genet 2012, 8(10).

9. Wei Q, Zhan XW, Zhong X, Liu YZ, Han YJ, Chen W, Li BS: A Bayesian framework for de novo mutation calling in parents-offspring trios. Bioinformatics 2015, 31(9):1375–1381.

10. Chang LC, Li B, Fang Z, Vrieze S, McGue M, Iacono WG, Tseng GC, Chen W: A computational method for genotype calling in family-based sequencing data. BMC Bioinformatics 2016, 17: 37.

11. Chen W, Li B, Zeng Z, Sanna S, Sidore C, Busonero F, Kang HM, Li Y, Abecasis GR: Genotype calling and haplotyping in parent-offspring trios. Genome Res 2013, 23(1):142–151.

12. Peng G, Fan Y, Palculict TB, Shen P, Ruteshouser EC, Chi AK, Davis RW, Huff V, Scharfe C, Wang W: Rare variant detection using family-based sequencing analysis. Proc Natl Acad Sci U S A 2013, 110(10):3985–3990.

13. Bishara A, Liu YL, Weng ZM, Kashef-Haghighi D, Newburger DE, West R, Sidow A, Batzoglou S: Read clouds uncover variation in complex regions of the human genome. Genome Res 2015, 25(10):1570–1580.

14. Zheng GXY, Lau BT, Schnall-Levin M, Jarosz M, Bell JM, Hindson CM, Kyriazopoulou-Panagiotopoulou S, Masquelier DA, Merrill L, Terry JM et al: Haplotyping germline and cancer genomes with high-throughput linked-read sequencing. Nat Biotechnol 2016, 34(3):303–+.

15. Edge P, Bafna V, Bansal V: HapCUT2: robust and accurate haplotype assembly for diverse sequencing technologies. Genome Res 2017, 27(5):801–812.

16. Conrad DF, Keebler JE, DePristo MA, Lindsay SJ, Zhang Y, Casals F, Idaghdour Y, Hartl CL, Torroja C, Garimella KV et al: Variation in genome-wide mutation rates within and between human families. Nature genetics 2011, 43(7):712–714.

17. Bishara A, Liu Y, Weng Z, Kashef-Haghighi D, Newburger DE, West R, Sidow A, Batzoglou S: Read clouds uncover variation in complex regions of the human genome. Genome Res 2015, 25(10):1570–1580.

